# GraSP Gene Targets to Hierarchically Infer Sub-Classes with CuttleNet

**DOI:** 10.1101/2024.04.26.591410

**Authors:** Samuel A. Budoff, Alon Poleg-Polsky

**Affiliations:** Department of Physiology & Biophysics, University of Colorado Anschutz Medical Center, Denver, CO 80204

## Abstract

This paper introduces a machine learning approach, GraSP, for retinal cell classification that addresses key challenges in spatial biology, alongside a novel neural network architecture, CuttleNet, tailored for class and subclass inference with incomplete datasets. We propose an innovative, unbiased gene selection method that utilizes simple neural networks for each target cell subclass, such that **Gra**dient **S**elected **P**redictors (GraSP) corresponding to gene importance are found for each. This approach significantly outperforms traditional machine learning techniques and expert-selected gene targets, reducing the necessary genes for classification from over 18k to 300 within the murine retina. Such reduction is crucial for advancing spatial biology, particularly in mapping retinal cell subclasses. Furthermore, our hierarchical architecture inspired by the organization of the cephalopod nervous system, CuttleNet, adeptly handles the pervasive issue of missing data in disjointed single-cell RNA sequencing datasets. CuttleNet operates by first classifying cell classes using consistently measured genes, then dynamically routing to subclass-specific subnetworks that leverage all available data for subclass classification. CuttleNet establishes a new standard in handling systematically missing data, offering substantial improvements over existing models in our targeted application.

## 1 Introduction

Spatial biology, building on insights from single-cell RNA sequencing (scRNAseq), aims to place defined cell classes and subclasses into their spatial context. A key challenge is the discrepancy between the number of genes measurable in scRNAseq and spatial sequencing (spSeq), which we address by reducing predictor genes from 18k to 300 [1; 2; 3].

The complex spatial arrangement of retinal cells serves as an ideal case study. Retinal cell classes such as photoreceptors, bipolar cells (BCs), amacrine cells (ACs), and ganglion cells (RGCs) overlap in gene expression but can be distinguished. Subclasses can be as simple and obvious as the rods and cones dividing the photoreceptor class [4] or as complex as the 45 RGC subclasses identified by scRNAseq [2]. Critically, all of these subclasses perform distinct computational roles that enable vision. However, historic methods have only made limited progress mapping these subclasses, spSeq techniques promise to overcome this but currently either lack XYZ resolution or can measure only hundreds of genes. Solving this challenge is critical to understanding how and why distinct spatial arrangements of these computational cells evolved.^2^

We bridge this gap by introducing: 1) A novel machine learning (ML) strategy for efficient gene target reduction, **Gra**dient **S**elected **P**redictors (GraSP), and 2) CuttleNet, a hierarchical NN architecture that addresses systematically missing observations in training datasets. GraSP enables a 60-fold reduction in gene targets needed to classify all subclasses in the murine retina, outperforming expert-chosen and standard ML tools. CuttleNet achieves state-of-the-art performance for hierarchical cell class and subclass inference, surpassing the best available supervised classifier [7].

### 1.1 Related Work

#### 1.1.1 Related Gene Selection Techniques

Various ML-based classifiers are employed in transcriptomic studies for dimensionality reduction, from classic methods like Random Forests (RF) [1; 8], to more complex tools based on graph theory [9; 10], and many techniques in between [11; 12]. A foundation model for scRNAseq data is even being developed, but to date is currently limited to human data and excludes retinal cells [13]. For the retina specifically, the Model-based Analysis of Single-cell Transcriptomics (MAST) framework was first applied to BCs [1] by identifying differentially expressed genes to account for cellular detection rates [14]. These gene targets were then used for classification with both simple thresholds as is traditional in histology, as well as RF [1]. However, MAST was replaced for the more nuanced RGCs and ACs [2; 3] in favor of a method of evaluating gene combinations based on the Area Under the precision-recall curve (AUCPR), emphasizing high precision to reduce false positives in cell-type labeling [15]. We show that performance relative to these classic and modern techniques can be improved upon with our GraSP algorithm and CuttleNet architecture.

Other Neural Network (NN) based strategies also treat observations as matrices of gene expression values, where each vector is associated with a cell subclass label. For example, Lin et al. [16] successfully used NNs for scRNAseq data clustering. However, their dataset contained 17k genes across 16 cell classes, relative to the 300 genes we use to classify 130 subclasses. We are limited to 300 genes due to technical limitations of the 10X’s Xenium platform at the time we developed GraSP.

#### 1.1.2 Related Hierarchical Networks

Hierarchical classification embodies the Coarse-to-Fine perceptual strategy. Projects like ThalNet [17] also draw inspiration from neuroanatomy. ThalNet uses static reading weights for temporal hierarchies, which is in contrast to the more anatomic approach we pursue here. Most related to our architecture, CellTICS leverages the Reactome gene-function map for interpretability in order to also distinguish cell classes and subclasses via supervised training with pre-labeled scRNAseq data [7]. This Reactome mapping identifies marker genes from biochemical pathways for classification. In contrast, CuttleNet employs generic sub-networks without prior biological annotation, mapping cell classes-to-subclasses by leveraging knowledge of dataset specific systemic problems like imputation. The dynamic routing we employ through tentacle-like sub-networks provides resistance to imputed values, overcoming observational discrepancies in real-world scRNAseq datasets. Our benchmarking experiments demonstrate CuttleNet’s superior performance and robustness compared to CellTICS in our limited application space.

## 2 Materials and Methods

### 2.1 Computational Resources

All experiments were conducted locally using consumer-grade components including an Intel i7-12700KF CPU, an NVIDIA RTX 3060 GPU with 12 GB of vRAM, and 128 GB of RAM (comprising four 32 GB Corsair CMK64GX5M2B5200C40 modules), all mounted on an MSI Z690-A Pro motherboard. The system operated on Ubuntu 22.04. All NNs and experiments were executed locally within a Python v3.9.16 Conda environment, utilizing Jupyter Notebook v6.5.4^*†*^ as an IDE or bash commands. The primary codebase was developed using NumPy v1.24.4^*†*^, pandas v1.5.2^*†*^, pyreadr v0.4.7^*‡*^, PyTorch v2.0.0^*†*^, scikit-learn v1.2.1^*†*^, and tqdm v4.64.1^*§*^. Additionally, comparisons with standard ML approaches and further data analysis and visualization were carried out in R 4.3.3^*‡*^, using the RStudio 2023.03.0 IDE^*¶*^, employing caret v6.0-90^*‡*^, e1071 v1.7-13^*‡*^, gbm v2.1.9^*‡*^, ggsignif v0.6.4^||^, matrixStats v1.0.0^*‡*^, multcomp v1.4-18^*‡*^, patchwork v1.1.3^||^, randomForest v4.7-1.1^*‡*^, rpart v4.1.23^*‡*^, Rtsne v0.16^*†*^, tidyverse v2.0.0^||^, and umap v0.2.10.0^||^. Licenses: *†* BSD-3-Clause License; *‡* GPL-2.0 License; *§* MIT License; *¶* AGPL-3.0 License; ||GPL-3.0 License. Code is available upon request prior to publication.

#### 2.1.1 Data Preparation and Preprocessing

Our dataset amalgamated data from 3 scRNAseq studies of retinal cells [1; 2; 3] available open source licensed as public data in Broad Single Cell Portal (studies SCP3, SCP509, and SCP919). Integration leveraged the log-transformed, median-normalized expression matrices computed by the original authors to account for batch effects between experiments and datasets. These studies originally contained 13166, 18222, and 17429 genes measured in 27500, 35700, and 32524 cells, respectively.

To enhance computational efficiency, we pre-processed the dataset by removing genes that did not contribute meaningful information, specifically those with zero expression or those consistently expressed across all cells. Using an empirically determined threshold of 0.5 for mean expression per subclass, we defined ubiquitously expressed genes and filtered them out. This process reduced the gene set by approximately ten-fold to 1642 informative genes. Missing genes in any dataset were assigned zero values to maintain matrix dimensions across samples, with the original dataset each cell was derived from stored as metadata to keep track of zero-imputations. The original subclass assignments from each study were standardized with a two-character prefix to ensure consistency across datasets, facilitating automated cell class assignments based on expert-defined rules.

The categorical cell class and subclass labels were numerically transformed utilizing Scikit-learn’s LabelEncoder in Python or R’s as.factor function. Prior to each model training, the dataset was shuffled (using a specified random state to ensure reproducibility) and divided into input features, **X**, and labels **y**. The shuffled dataset was finally split into training (80%) and test (20%) sets.

### 2.2 Evaluation Criteria And Cross Validation

For all ML methods evaluated, we define a subclass as satisfactorily classified if they were done so with at least a value of 0.9 for both the True Positive Rate (TPR) and Precision (Prec) computed from confusion matrices on the testing dataset. For comparisons of specific classes or subclasses the F1 statistic was evaluated, again computed from the testing set. To ensure the robustness of our findings, all models and experiments included in this work were repeated five times, each with a different random seed to initialize the shuffling and subsequent calculations. These replicates allowed ANOVA testing for statistically significant differences in the models evaluated in an experiment, followed-by post-hoc Tukey HSD testing of all pairs of models/conditions, with appropriate p-value adjustment. For all plots, * indicates p<.05, ** p<.01, and *** p<.001. Standard box-plots generated in R extend whisker to largest value *≤* 1.5*\**IQR.

### 2.3 Traditional ML-Based Gene Reduction

The MAST [14] and AUCPR [15] methods were used to propose published sets of genes for classifying the 130 cells in the retina, so we evaluated these models’ outputs [1; 2; 3]. The intended use of these genes in mapping experiments is to evaluate presence/absence to classify a given subclass. In practice, many laboratory techniques require a threshold to define said presence/absence. We simulated this for a set of RGCs’ AUCPR predictors by evaluating the F1 performance with the following thresholds for gene presence/absence: **0** expression detected, **50th** greater than the 50th percentile, **75th** greater than 75th percentile, and **90th** greater than the 90th percentile.

We evaluated the overall ability for MAST and AUCPR to classify retinal subclasses by training NNs with all of the MAST proposed BC predictors[1], the AUCPR proposed RGC [2] and AC [3] predictors, and the combination of these. The NNs were of identical architecture and training protocol as the GraSP NNs described below. We also employed several standard ML models to suggest sets of 300 genes from our scRNAseq dataset to determine how many subclasses could be accurately classified with only this reduced set of predictors by said model. The classic models we investigated included: Decision Trees (DT) [18], RF [19], Gradient Boosting Machine (GBM) [20], and Support Vector Machines (SVM) [21]. Each model was first trained on the full training split. Post training, the importance of each gene was assessed to extract the top 300 genes. For DT, RF and GBM, Gini impurity was used to define the importance of a given gene. For the SVM, importance was determined by the absolute values of the coefficients in the linear kernel. These ML models were implemented in R from standard libraries. After proposing a set of 300 genes, the training set was subset accordingly and each model was retrained. The performance of these models was evaluated by the number of subclasses classified above 0.9 TPR and Prec.

### 2.4 GraSP Gene Reduction

#### 2.4.1 An Ensemble of Specialists

The core of our methodology involved training an ensemble of simple NNs; *f*_w_(**x**) with weights **w** mapping the input gene expression vector **x** to the output class probabilities. Each NN is identical in architecture but trained to become a specialist for a specific cell subclass, *c* ∈ {1, 2, …, *C*}, by training on a purposefully biased training dataset. Let 𝒟 be the training dataset, where 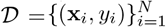, with **x**_*i*_ *∈* ℝ^*d*^ being the input gene expression vector for the *i*-th sample and *y*_*i*_ ∈ {1, 2, …, *C*} being the corresponding class label. *𝒟*_*c*_ is the biased training dataset for class *c*. These training sets *D*_*c*_ are automatically formed by combining all observations of target subclass *c* from 𝒟, totalling *N*_*c*_ samples, with an equal number, *N*_*c*_, of randomly sampled distractors from the remaining classes, *d* ∈ {1, 2, …, *C*} − {*c*}. Formally, 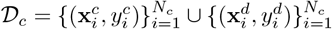.

These NNs were fully connected and had an input layer where each node represented an available predictor gene (1642 input nodes), an output layer with each node representing a subclass label (130 output nodes), and a single hidden layer with twice as many nodes as subclasses (260 nodes). ReLU non-linearities were used on the input and hidden layers, with log-softmax used on the output. Cross-entropy loss and the Adam optimizer with learning rate = 0.001 were used for training 100 epochs on batches of 32 observations.

Formally, let ℒ (*f*_w_(**x**), *y*) be the loss function, which measures the discrepancy between the predicted class probabilities *f*_w_(**x**) and the true class label *y*. Each NN, 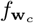, becomes specialized for class *c* by training on the biased training dataset 𝒟_*c*_ by minimizing the loss function over the biased dataset:

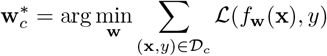

#### 2.4.2 Gradient Selected Predictors

To identify the most relevant genes for cell subclass classification, we devised a gradient-based feature importance statistic. After training each specialist NN with the respective biased datasets, the gradients of the loss with respect to the input predictors, which in our case, were gene expression levels, were calculated. Formally, for each sample (**x**, *y*) *∈* 𝒟_*c*_, we computed the gradients of the loss function with respect to the input gene expression vector **x**:

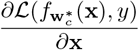

The mean absolute value of these gradients across the batch dimension was used as a measure of each gene’s importance in classification. To calculate the mean absolute value of the gradients across all samples in 𝒟_*c*_ for each gene *j* ∈ {1, 2, …, *d*}:

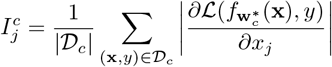

The gene importance scores 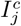 represent the sensitivity of the loss function to changes in the expression level of gene *j* for classifying class *c*. Higher values indicate that the gene is more important for discriminating class *c* from other classes. This method captures the network’s sensitivity to changes in gene expression levels, ranking the genes’ influence for classification of *c*.

The feature importance calculation was performed for each subclass-specialized network, providing a rank order in descending order according to their importance scores 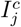 for each class *c*. The rank order of all predictors relative to all subclasses was then compiled into a list by iterating through each subclass’ individual list and storing novel genes *j* as they were encountered. A final NN model with the above architecture (except only 300 input nodes) was trained using the selected gene set. This model was evaluated on the test set to assess its classification performance.

#### 2.4.3 Ablation Study

ML models may take advantage of the zero-imputation we used to maintain matrix dimensionality. On the other-hand, discarding such predictors is not desirable when they are truly informative. To test this we performed an ablation study by comparing the classification performance of specific classes and subclasses predicted by a given model trained with or without zero-imputed genes. A paired T-test was used to test for changes caused by this manipulation.

### 2.5 CuttleNet - Hierarchical Network Architecture

To avoid such cheating without deleting real data, we developed a novel two-tier NN, CuttleNet, designed to hierarchically classify cell classes followed by cell subclasses. CuttleNet consists of a primary class classifier NN and multiple subclass classifier NNs, the general architecture of each is identical to the fully-connected network described above for GraSP, see **Fig. 3A**.

#### Class Classifier

The first stage predicts cell classes with input consisting of data common to all studies, and the output layer size corresponds to the number of classes. Non-linearity is introduced via ReLU activation functions, and a softmax layer generates a probability distribution over classes.

#### Subclass Classifiers

Each subclass classifier, stored in a PyTorch ModuleDict for dynamic selection, is tasked with identifying subclasses within a predicted class. These networks take concatenated inputs of class-specific gene expression data and the class classifier’s output. They are uniquely configured to output predictions for the number of subclasses they represent, based on a predefined class-to-subclass mapping. The class-specific genes are simply those free of zero-imputation and thus correct the main methodological compromises in the dataset merging.

#### 2.5.1 Benchmarking

To assess the performance of CuttleNet relative to the best available model for this hierarchical task, 1 layer and 5 layer CellTICS models were trained with and without access to imputed data. We first optimized the CuttleNet performance at classifying the maximum number of subclasses via a systematic grid search varying epochs, early stopping, and L1 regularization: **Number of Epochs:** Tested across 10, 25, 50, 100 epochs to assess the impact on model convergence. **Early Stopping:** Implemented with 0, 5, 10 epochs to prevent overfitting. **L1 Regularization:** Applied with 0.0, 0.001, 0.005, 0.01, aiding in feature selection and robustness. Each hyperparameter combination was evaluated across multiple seeds (18 to 108, in increments of 18) for robustness and reproducibility. The model’s performance was assessed on a test set following training with the specified configurations.

## 3 Results

### 3.1 Gene Selection

spSeq promises to comprehensively map retinal cells, but to facilitate this it is necessary to determine a much smaller set of genes capable of effectively and reliably clustering cell subclasses. Traditionally, biologists rely on expert selected “gold-standard” targets to classify cells in their native spatial positions [4; 5; 22; 23; 24; 25; 26]. scRNAseq allowed for experts to propose not only novel cell subclasses, but also predictors identified by much more powerful statistical techniques, such as MAST [1; 14] or AUCPR [2; 15]. These small sets of genes, relevant for traditional experimental mapping methods, are typically limited to *≤* 3 per subclass where a binary assessment of presence or absence is used for classification. While spSeq cannot make use of the *>* 10^4^ genes available to scRNAseq, it can use many more than 3 genes at a time, and in a more nuanced manner than presence or absence. To test if published gene sets proposed by statistical methods including PCA and AUCPR [2; 15] could be useful to spSeq we evaluated the proposals of 10 RGCs by setting a variety of thresholds, as one would do in an experimental setting, and performed classification. The F1 performance of such small gene sets indicated that neither expert-chosen genes nor traditional thresholding strategies can be relied upon for spSeq, as shown in **Fig. 1A** and **Sup. Fig. A.5**. ^3^.

**Figure 1.**
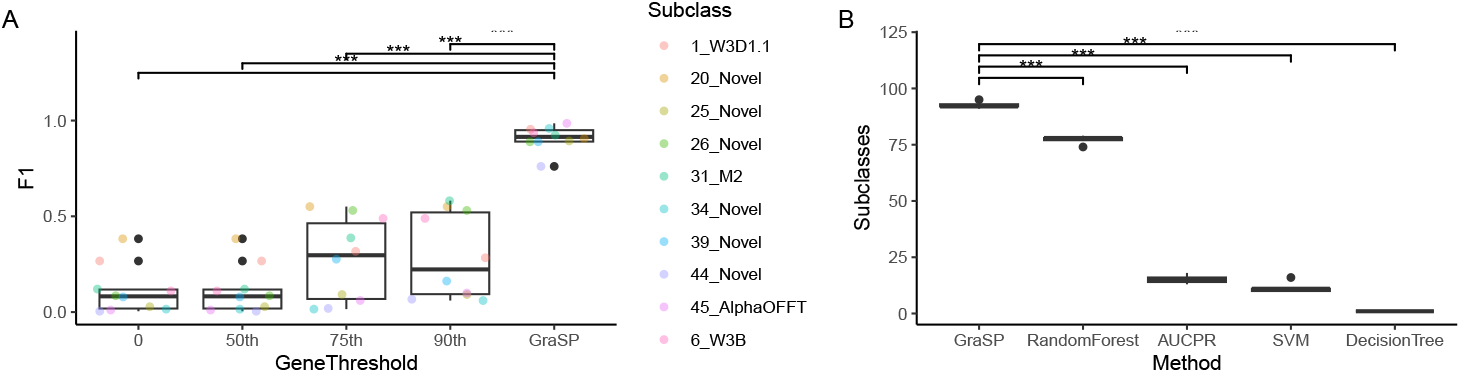
GraSP performance relative to: **A)** Published gene targets proposed by AUCPR with classification thresholds simulating laboratory practice; **B)** Traditional ML methods used for selecting 300 genes as well as a NN trained on the published AUCPR set of genes for classifying all 130 retinal subclasses with classification defined by subclass performance above 90% TPR and Prec.

To identify gene sets more purposefully designed for spSeq we explored traditional ML methods for dimensionality reduction. The most commonly used embedding tools, like UMAP, t-SNE and PCA projections do not explicitly preserve the importance of individual predictors so were not relevant. Methods which do, such as RF and SVM, however, did not perform as well as we expected, or simply failed to converge, as was the case for genetic evolutionary algorithms. While exploring, we observed that very simple feed-forward NNs could be trained via back-propagation to predict subclasses in a supervised manner with superior performance to these methods. This inspired us to interrogate the gradients computed by these NNs from a given subclass back to its predictors. In so doing we devised an importance statistic that could rank order these genes for a given subclass. As our dataset is skewed over 4 orders of magnitude (**Sup. Fig. A.4**) we decided to create biased training datasets for each subclass, to train an ensemble of NNs where each was a specialist at its respective subclass. Computing and combining the predictors selected by the gradients’ importance score produced our GraSP algorithm for gene selection.

GraSP significantly outperformed published predictor gene sets, **Fig. 1A**, ANOVA results indicate a significant difference in F1 scores among different gene thresholds (*F* (4, 45) = 42.07, *p* = 0). Post-hoc Tukey’s HSD test revealed that the network chosen gene thresholds significantly outperformed the human chosen thresholds, with *p* = 0 for all comparisons. Practically, the GraSP chosen genes demonstrate superior performance, with average F1 performance at least 60% better than the best thresholded published gene sets. Even when all predicted genes from AUCPR, and MAST are used to train NNs identically to the final NN containing GraSP predictors, they fail to satisfactorily classify the potential set of subclasses **Sup. Fig. A.5**. This highlights the efficacy of the GraSP approach over published retinal classification schemes in the context of spSeq.

To evaluate our novel algorithm relative to traditional ML methods, we defined subclasses as being correctly classified with the selected predictors if they were classified with a TPR and Precision both above 90%. Using the GraSP, DT, RF, GBM, and SVM we asked how many subclasses could be classified with 300 genes. We also combined all AUCPR and MAST published predictors for the retina and used them to train a NN identically to the final network proposed by GraSP. ANOVA results indicate a significant difference in the number of subclasses classified by the different methods (*F* (4, 20) = 2770, *p* = 0). Post-hoc Tukey’s HSD test revealed that GraSP significantly outperformed each method with *p* = 0. Practically, GraSP demonstrated a substantial advantage in classifying subclasses. Specifically, GraSP identified an average of 15.4 more subclasses than RandomForest, 81.0 more than SVM, 91.6 more than DecisionTree, and 77.4 more than the AUCPR and MAST published predictors - highlighting its superior performance compared to traditional machine learning methods, see **Fig. 1B**. For completeness, we also evaluated the individual sets of MAST, AUCPR, and historical expert selected predictors for the respective subclasses they targeted by again using them to train NNs and evaluated them on the percent of applicable subclasses classified above the 0.9 TPR and Prec threshold, **Sup. Fig. A.5**. In this comparison as well GraSP significantly outperformed them, *F* (4, 20) = 2120, *p* = 0 Tukey HSD *p* = 0 against each published predictor set and their combination.

GraSP is significantly faster than traditional methods, selecting genes and training the final model in about 8 minutes, similar to DT, but much faster than the 2 hours for RF, and 6 hours for SVM. The Gradient Boosting Machine was also explored as a traditional method, but with the full list of available genes it ran for 36 hours before it was abandoned due to inefficiency - The completed iteration classified only 39 subclasses. These efficiency differences are because NNs train on GPUs, whereas these traditional methods are sequential and as implemented in R train on the CPU. GPU clusters or simply devices with more vRAM would train GraSP even faster.

Finally, we clustered the GraSP genes using both UMAP and t-SNE projections to ensure we were not being misled by our target statistic. Encouragingly, we reproduced published clusters with both techniques - validating that the separation of clusters is preserved regardless of embedding algorithm as shown in **Fig. 2** and **Sup. Fig. A.6**. This observation indicates the robustness of our novel method for preserving the practical outcome of our dimensionality reduction, ultimately validating that we could reduce the number of genes needed for clustering by 60 fold.

**Figure 2.**
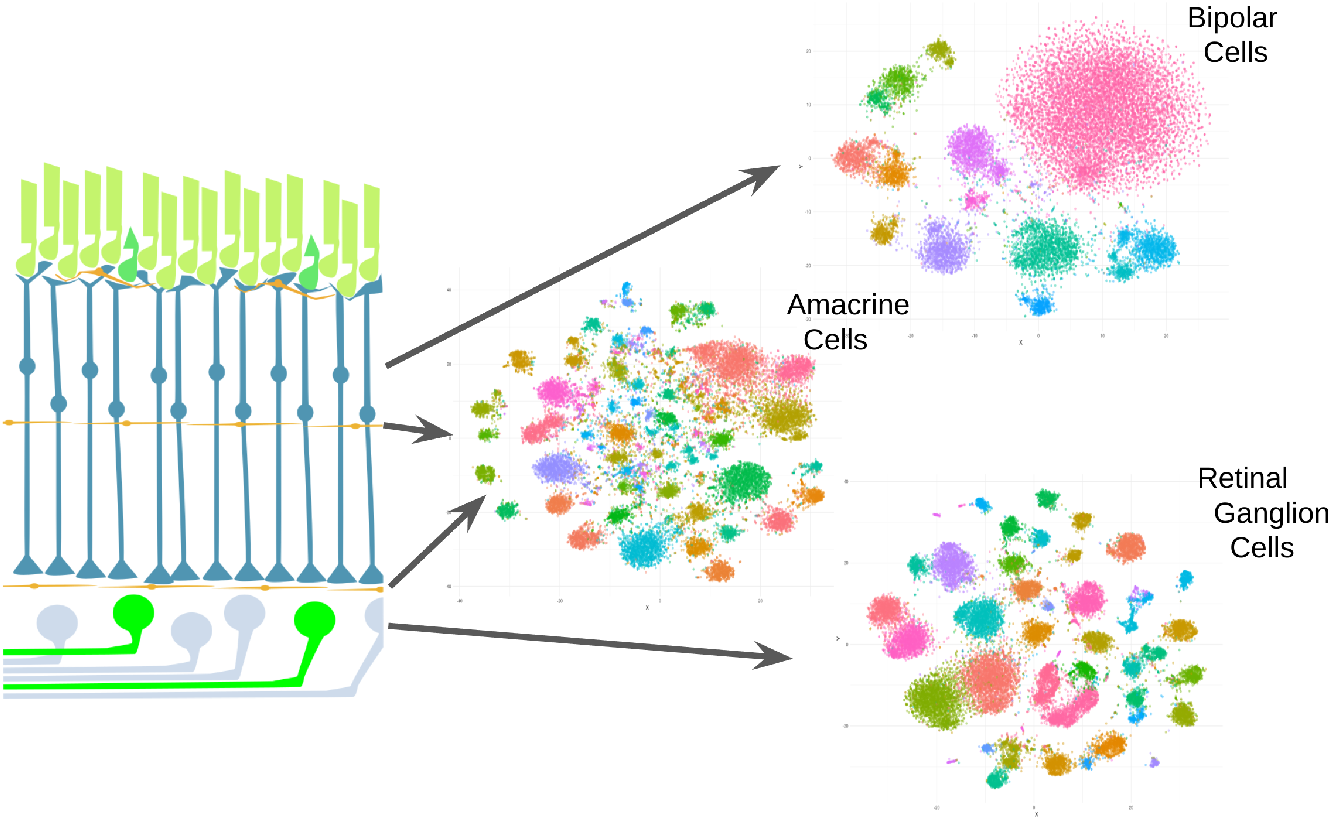
UMAP projection of GraSP minimized retinal cell gene vectors for the indicated classes with spatial location of these classes indicated by the diagram of a retinal cross section.

**Figure 3.**
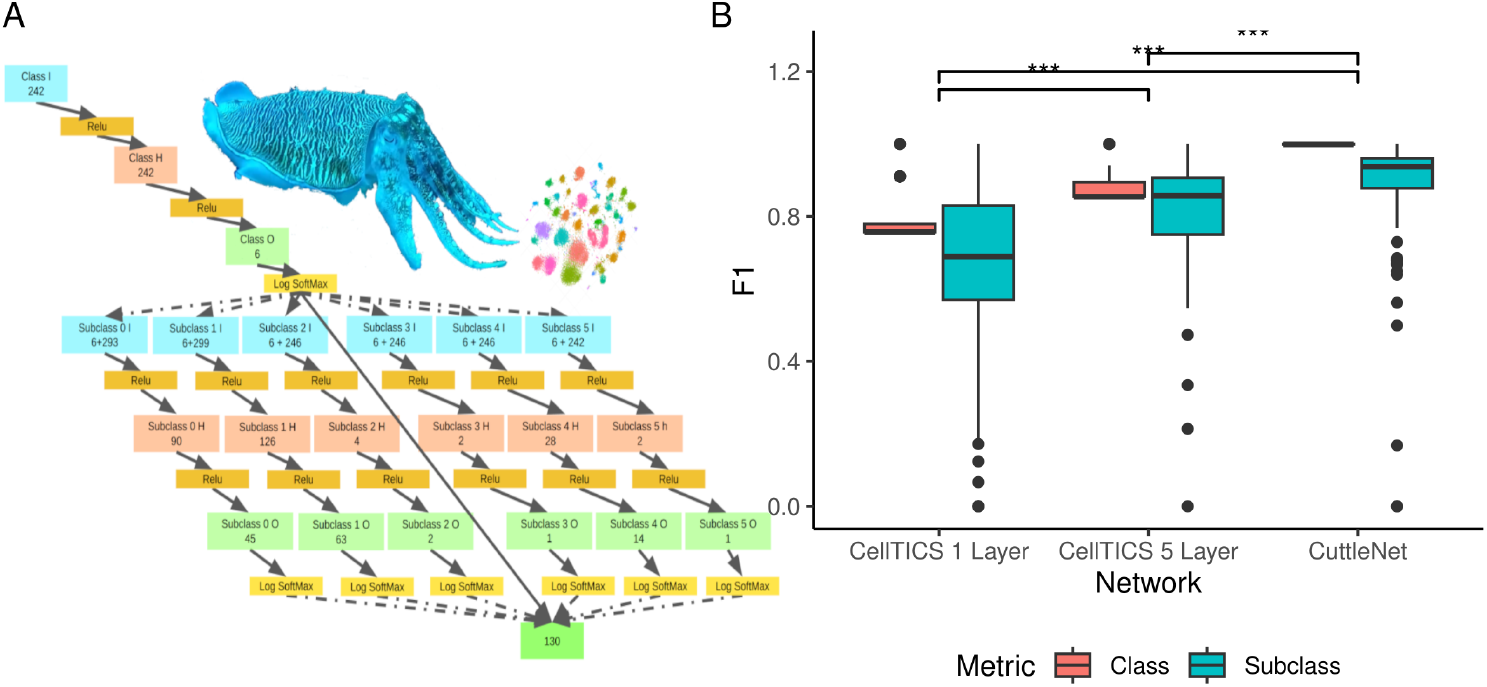
**A)** CuttleNet diagram, with Carl the Cuttlefish grasping at cell clusters. Box colors indicate: Blue = Input Layer, Orange = ReLU, Salmon = Hidden Layer, Light Green = Inner Output Layer, Green = Final Output Layer. Number of nodes indicated in each layer with 6 + *g* in the subclass’ input tentacle indicates the concatenation of the class output with the *g* genes available to that layer. Dashed arrows indicate conditionally engaged connections. **B)** Benchmarking CuttleNet vs. CellTICS.

### 3.2 CuttleNet

A worrisome compromise we introduced in creating a pan-retinal cell expression matrix from independent studies of cell classes was the imputation of genes not measured between studies. That is, if a gene was found to be sufficiently variable to pass our initial culling of one expression matrix, it is not guaranteed to have been measured in the other scRNAseq studies. Instead of discarding such candidate genes, we imputed values of zero for studies which did not include them. We suspected that some of the performance of our NNs might stem from this decision. To test this, we performed an ablation study by removing all genes with imputed 0s from our GraSP subset expression matrix and trained a version of our NN. This caused a slight but significant performance drop using a paired T-test of our subclass classification F1 score, *t*(153299) = 19.3, *p* = 2 *×* 10^−16^. The practical significance in this case is small, but it gave additional credence to our concern about the impact of imputed values, **Table 1**. Thus, we devised a hierarchical modification to our NN, leveraging biologic design principles in which separate NNs acted as sub-networks responsible for either classes, or a specific set of subclasses. Such an architecture, like that found in a cuttlefish’s nervous system, allows for quasi-independence of networks, where subregions of the network receive from upstream regions only the information necessary for their task, but all of the sub-networks ultimately act together to perform complex behaviors. The specific information entering a given sub-network was the set of available genes free of imputed 0s, and samples predicted to be the respective class. The ultimate output was an aggregation of the activated subclass-specialized sub-network’s predictions (**Fig. 3A**).

**Table 1:**
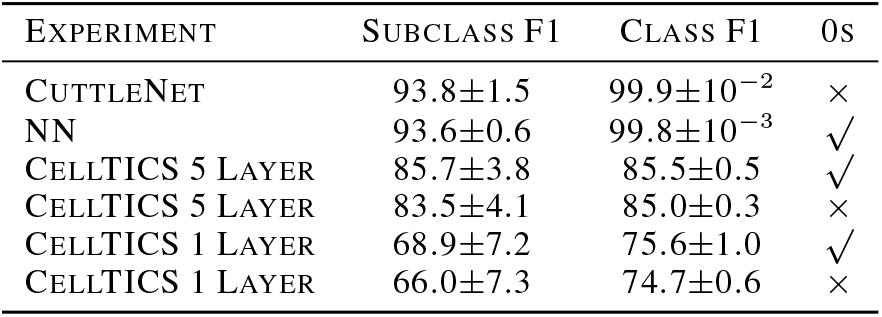
Ablation Study: Median *±* standard deviations F1 scores. 0s shows if model uses imputations.

We finally benchmarked our networks against CellTICS, the most cutting-edge hierarchical NN model for simultaneous cell class and subclass classification [7]. Appreciating the extensive benchmarking of CellTICS against previously leading methods, and its superior performance to them, we deemed it the best reference for evaluating CuttleNet. To fully compare it, we trained it with versions of our retina expression matrix both with and without imputed values. Furthermore, we trained both the 1 Layer architecture, which is most similar to our own, and the developer’s recommended 5 Layer network (**Sup. Fig. A.7**). To ensure robust comparisons, we re-trained and evaluated CellTICS five times. Before benchmarking, we performed a comprehensive grid search of CuttleNet’s training hyperparameters to optimize it as CellTICS had been (**Sup. Fig. A.8**).^4^

CuttleNet demonstrated significant superiority over CellTICS for our specific application. We performed ANOVAs and post-hoc Tukey tests on the class and subclass F1 scores for CuttleNet and the CellTICS models with 1 layer and the developer optimized 5 layers. In both the class and subclass-focused ANOVA, we rejected the null hypothesis as expected from **Fig. 3B**, with *f* (2, 372) = 939.3, *p* = 0 and *f* (2, 357) = 54.13, *p* = 0, respectively. It should come as no surprise based on **Fig. 3B** that post-hoc testing also rejected the null hypothesis for each of our models against CellTICS with classes and subclasses with *p* = 0 and *p* = .009 for the 5 Layer for subclasses. In comparing CuttleNet with the simpler NN, in a separate ANOVA, it is worth highlighting that for class classification performance CuttleNet is significantly better (*p* = .006), as with subclass performance (*p* = .012). That said, the practical difference in performance between the models is small, with a Cohen’s *d* = 0.228. This demonstrates that CuttleNet does not sacrifice power as a result of its added complexity. By contrast, CuttleNet is a superior model for inference because it selectively ignores imputed values, and thus solves a real-world problem facing researchers who have had to merge datasets in this very common manner. Our ablation studies highlight (**Table 1**) how models might use the 0s as “obvious” class markers to cheat. In our case, the practical difference of this cheating is small, but on other datasets, it could have a meaningful impact.

## 4 Discussion

The GraSP method presents a novel approach to feature selection in the context of transcriptomic data analysis. This method addresses a crucial challenge in spSeq studies, where the number of measurable targets is constrained by technical limitations, and thus predictor targets must be minimzed while maintining their ability to discriminate subtly different classes. In our study, the gene set was reduced from approximately 18000 to just 300, without compromising classification performance. This dramatic reduction not only demonstrates the efficiency of our neural network strategy but also lays the foundation for future research where such constraints are commonplace. As spSeq technologies continue to advance, researchers may still opt for minimal classification panels due to cost and time considerations, further underscoring the relevance of our approach.

The core of the GraSP method lies in the calculation of the gene importance score 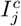, which quantifies the influence of each gene *j* on the classification performance for a specific target class *c*. By computing the mean absolute value of the gradients of the loss function with respect to gene expression levels, GraSP captures the sensitivity of the neural network’s predictions to differences in gene expression. This approach effectively ranks the genes according to their discriminative power.

Unlike traditional filter, wrapper, or embedded ML methods, GraSP employs a novel approach that does not neatly fit into these categories. GraSP shares some conceptual similarities with embedded methods in using neural networks to evaluate feature importance, or ensemble methods like RF where multiple models are created and contribute to predictions. Unlike these ML tools, GraSP trains an ensemble of specialized neural networks, each focused on a particular target class due to the biased training datasets and then computes gradients of the loss function to provide class-specific feature rankings for each. The novelty thus lies in compiling these rankings into a final list by iterating through each class and adding novel features, prioritizing the most discriminative genes across all classes. This approach, coupled with the use of biased datasets tailored to target classes, distinguishes GraSP from traditional ensemble techniques and embedded methods, making it a unique contribution to feature selection methodologies.

A **limitation** of the current implementation of GraSP is the unoptimized search strategy, thus the algorithm lends itself to further algorithmic improvements. For instance, a simple extension could involve drawing from *I*^*c*^ until a specified classification threshold is met for each class *c*, potentially enhancing the efficiency and interpretability of the gene selection process. Additionally, alternative functions, such as the square of the gradients, could be explored to capture different aspects of gene importance. This is an important future direction necessary to generalize our contribution.

A notable contribution of this work is the introduction of CuttleNet to address the challenge of imputed values in merged datasets; a common issue in many real-world scenarios where data is incomplete or derived from multiple sources. By segregating and processing class-specific information through separate sub-networks, CuttleNet demonstrates robustness in the presence of imputed data, enhancing its applicability to practical spSeq studies. The success of CuttleNet underscores the value of biologically-inspired architectures in addressing complex machine learning challenges.

It is important to note that CellTICS is a state-of-the-art hierarchical model designed to predict classes and subclasses from full scRNAseq studies, not the minimal constraints imposed by spSeq. Thus, it is not fair to take away from our results that CuttleNet is generally more powerful than CellTICS. On the other hand, CuttleNet is a unique model in that it is specifically designed to account for imputed values, which is a common challenge outside of the context of transcriptomic studies. Thus, future work could extend CuttleNet to other application spaces to evaluate how it can be used to handle this common dataset merging problem. Additionally, CuttleNet allows for each sub-network to have different internal architectures than the simple single hidden layer we here evaluated rigorously. Future work leveraging the general architecture to overcome data-cleaning issues can be easily imagined, as can be implementations with more than two hierarchical levels, which we.

### 4.1 Conclusion

In summary, the GraSP method and CuttleNet architecture represent novel contributions to the field of spatial biology and feature selection. By effectively reducing the gene set required for accurate cell subclass classification and addressing real-world data challenges, our work paves the way for more efficient and practical applications of spatial transcriptomics in biological research.

## Acknowledgements

We would like to thank Drs. Joel Zylberberg, Yaning Liu, Joshua French, and Erin Austin for friendly reviews of an early draft of this manuscript. We would also like to acknowledge that LLMs were used to support the writing of this manuscript and codebase. Specifically, the OpenAI chatGPT models using GPT-3.5, 4, and 4o, as well as Anthropic’s Claude with the Clause 2 and Sonnet architectures. The primary use of these models was to replace Stack Overflow from the author SB’s workflow for function and syntax look-up. Human written drafts of this manuscript were used to prompt these models for clarifying suggestions, as well as to simulate reviewers. Perplexity.ai was used to augment literature review in addition to google scholar, bioRxiv and, similar standard repositories of academic literature.

## A Appendix

### A.1 Subclass Proportions in Full Dataset

**Figure A.4:**
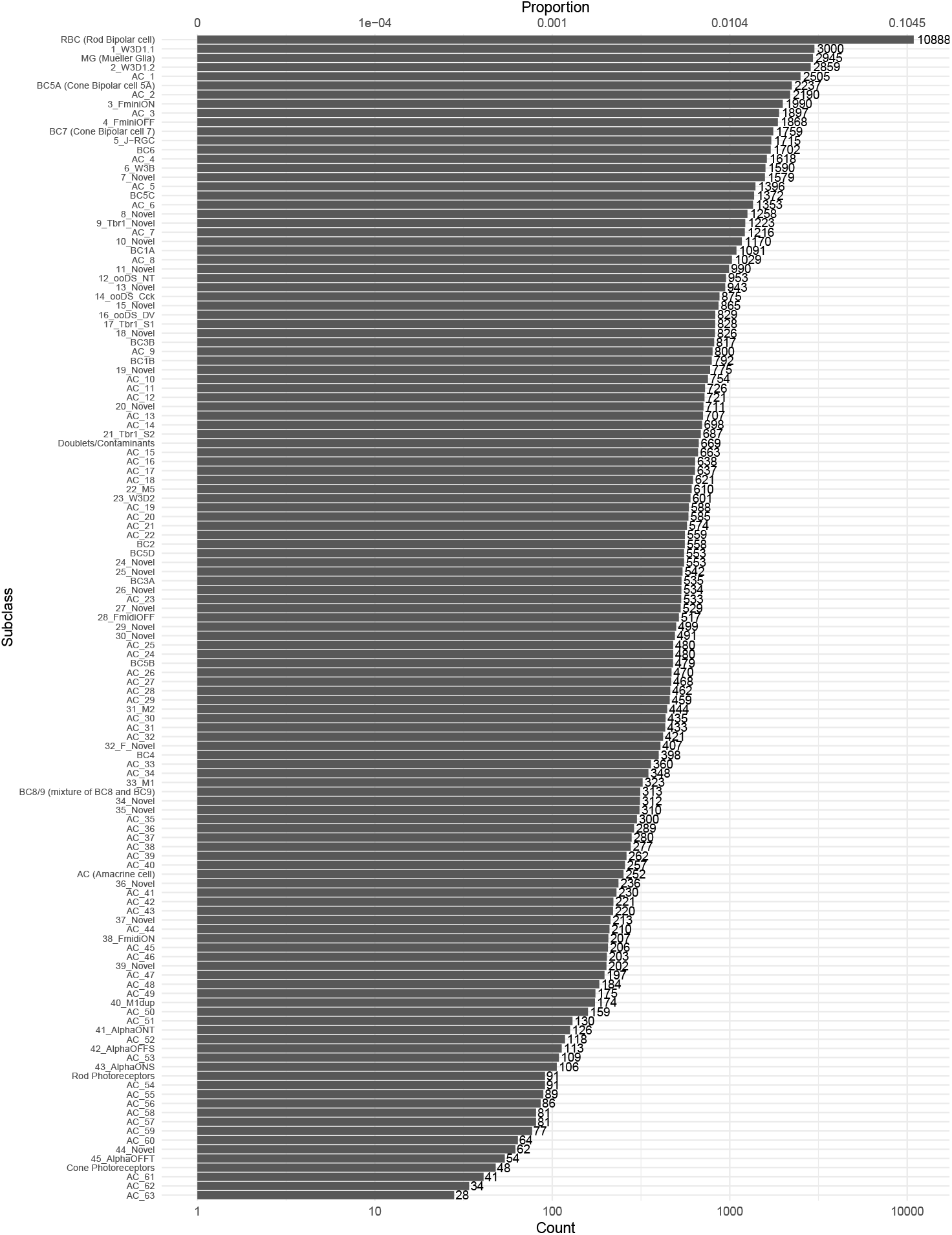
Counts and proportions of each cell subclass on a log scale demonstrating the dramatic imbalance in our targets.

### A.2 AUCPR & MAST Further Comparison

**Figure A.5:**
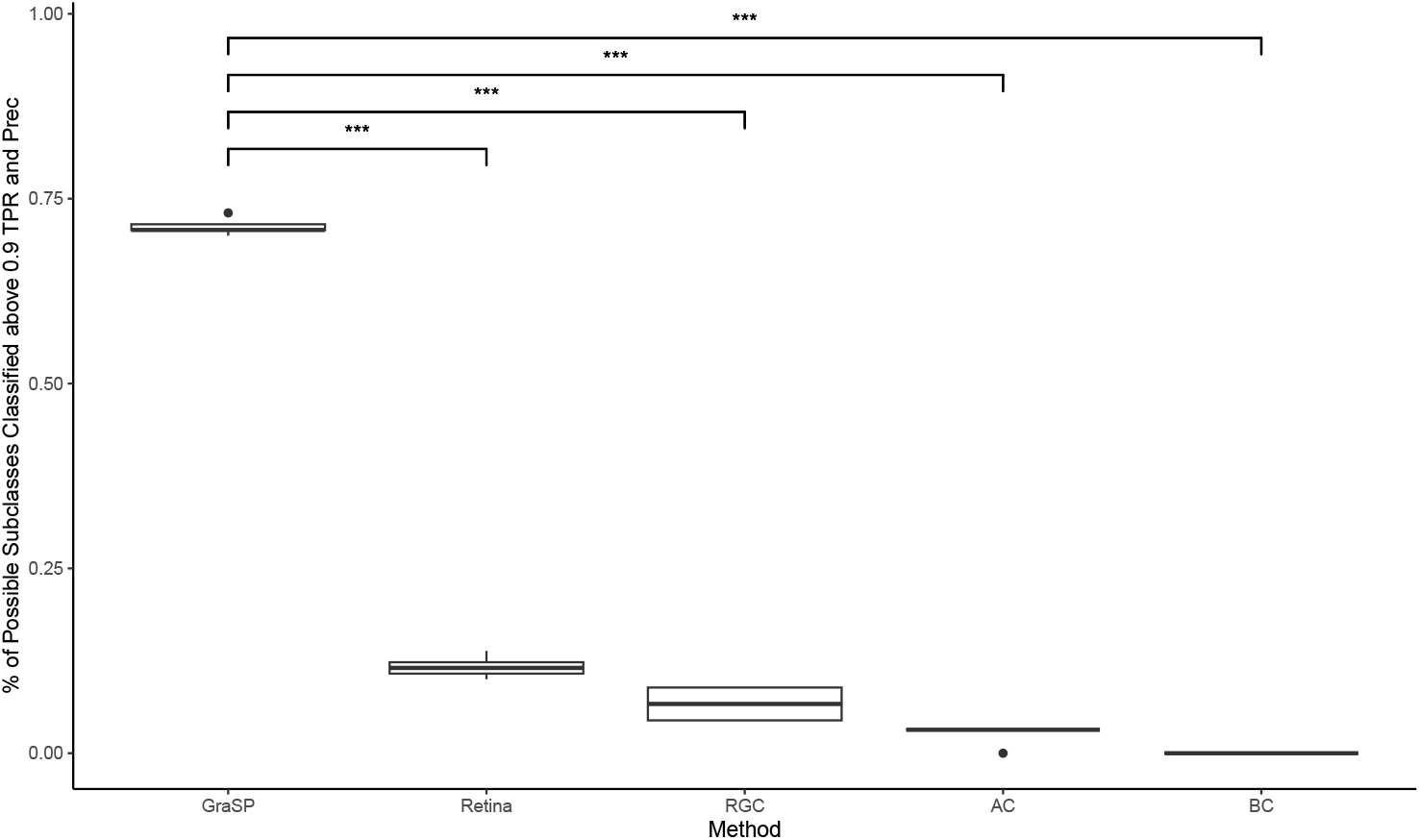
Percent of cell subclasses classified above 0.9 TPR and Prec out of the total number of subclasses targeted in the original study. BC used the MAST [14] method to classify 17 subtypes [1], RGC and AC used the AUCPR method [15] to classify 45 [2] and 63 [3] subtypes, respectively, Retina combined these independent sets to classify all retinal cells. GraSP also targeted all retinal cells. All classifiers were simple NNs with identical architecture and training other than the set of genes in their training data and thus input nodes.

### A.3 UMAP & t-SNE Clusters

**Figure A.6:**
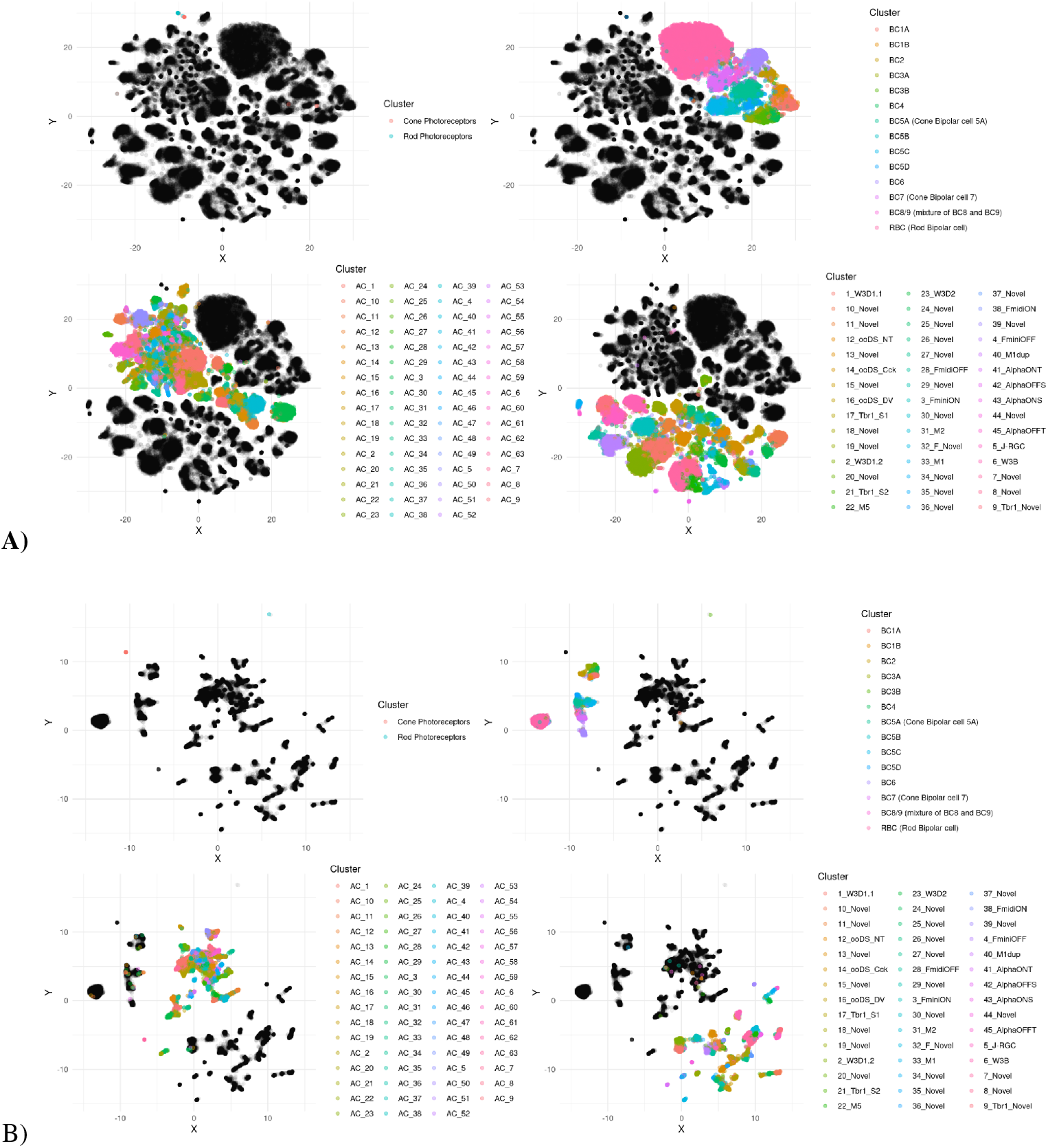
Projections of cells in test set with GraSP predicted genes only, with colors labeling respective subclasses within an expert defined class **A)** UMAP projection, **B)** t-SNE projection. Regardless of choice of projection in this case, we observed classes cluster close together, and at the same time subclass clusters are not mixed. Note that this is not generally going to occur with these methods but the fact that it occurs here is good evidence data separability is preserved. **Fig. 2** has the same subclass coloring as shown here, but the legend was removed for legibility in the diagramatic context. Note, that **Fig. 2** was a re-implementation of the UMAP clustering shown here, with data limited to cells within a retinal cell class in order to better reveal the homogeneity of clusters and their separation, which is difficult to shown in a static diagram like this.

### A.4 Grid Search Technical Details

This search was a deliberate exploration to pinpoint a balance between training duration, regularization, and early stopping criteria that would yield the best performance for both class and subclass predictions. The optimal parameters emerged at 25 epochs of training, without early stopping, and with a regularization strength of 0.001 **Sup. Fig. A.8**). This configuration not only maximized the subclass performance—with diminishing returns observed beyond—but in general, the grid-search revealed that this architecture rapidly converged on high-quality class predictions, achieving near-perfect accuracy before 10 epochs (**Sup. Fig. A.7**)). This aptitude for class identification may be due to the relatively simple task.

The nuanced role of the regularization parameter in our CuttleNet’s performance became evident when examining the longevity of class classification accuracy and subclass differentiation over extended training periods. The optimal regularization strength was found to be 0.001, striking a balance that mitigates overfitting while preserving the network’s ability to generalize. This was an important finding, as it allowed the network to maintain a high level of class classification accuracy, even with prolonged training epochs. Specifically, we observed that without regularization, the F1 scores not only dropped for all cell classes, but the variability between training replicates dramatically increased proportional to training epochs. Conversely, as regularization increased, the network’s capacity to distinguish between the more granular subclass categories diminished. This inverse relationship between regularization strength and subclass accuracy underscores the intricacies of model tuning for our hierarchical architecture (**Sup. Fig. A.7**)).

Note that earlier experiments when we were designing GraSP also involved a similar grid-search strategy, including over several other hyperparameters such as hidden layers and custom-loss functions, **Sup. Fig. A.9**). The performance differences between these permutations were all negligible, so for all work in this study the un-optimized, simple NN described in the methods was used. We are including this grid-search for completeness, however, in this case it does not impact our findings one way or another.

**Figure A.7:**
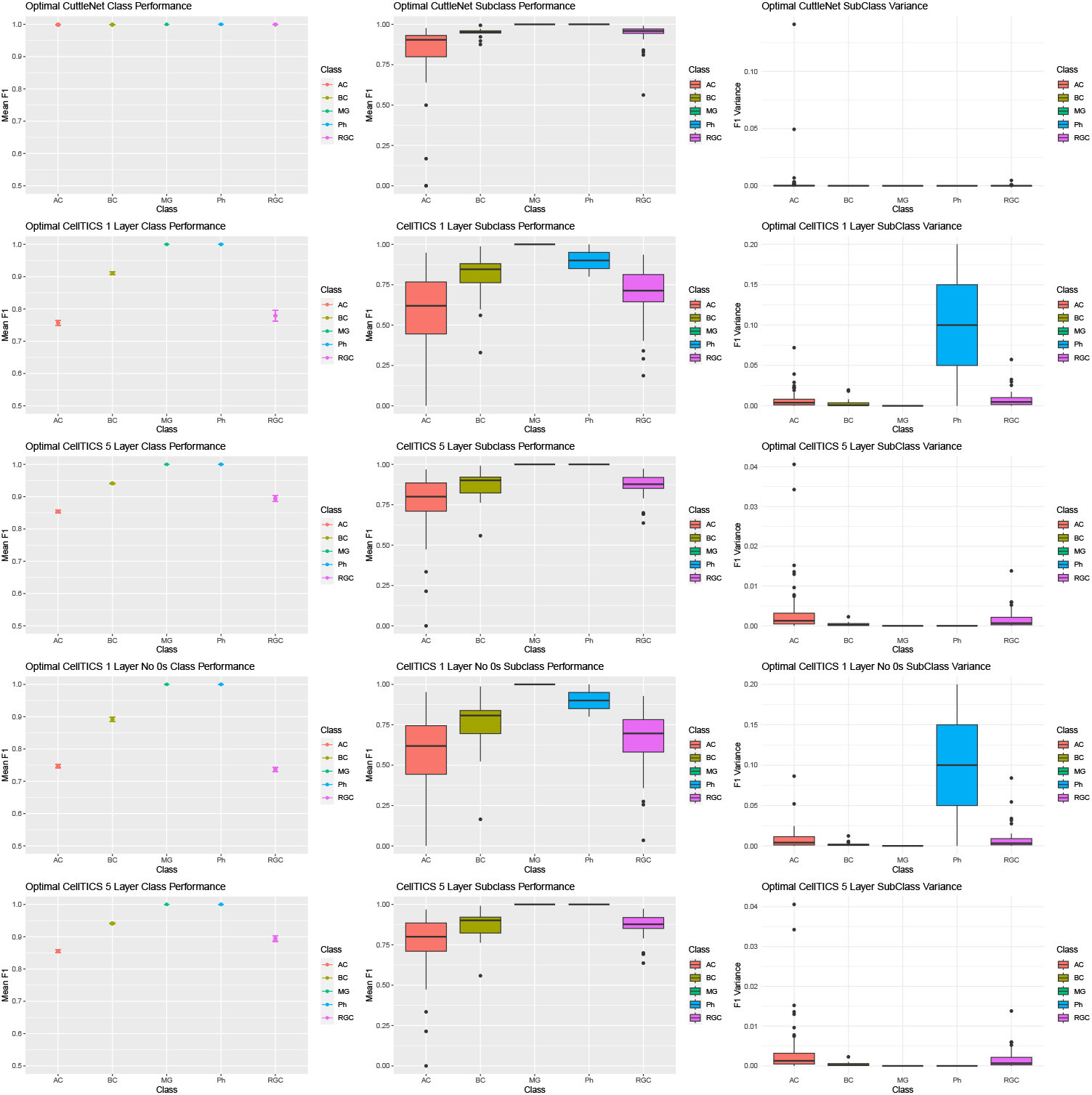
Optimal versions of CuttleNet found after grid search relative to the performance of the CellTICS models trained with 1 or 5 layers and access to the imputed values or not in the GraSP selected genes. Performance is shown for class and subclass classification performance as well as the variance over the 5 validation shuffles.

**Figure A.8:**
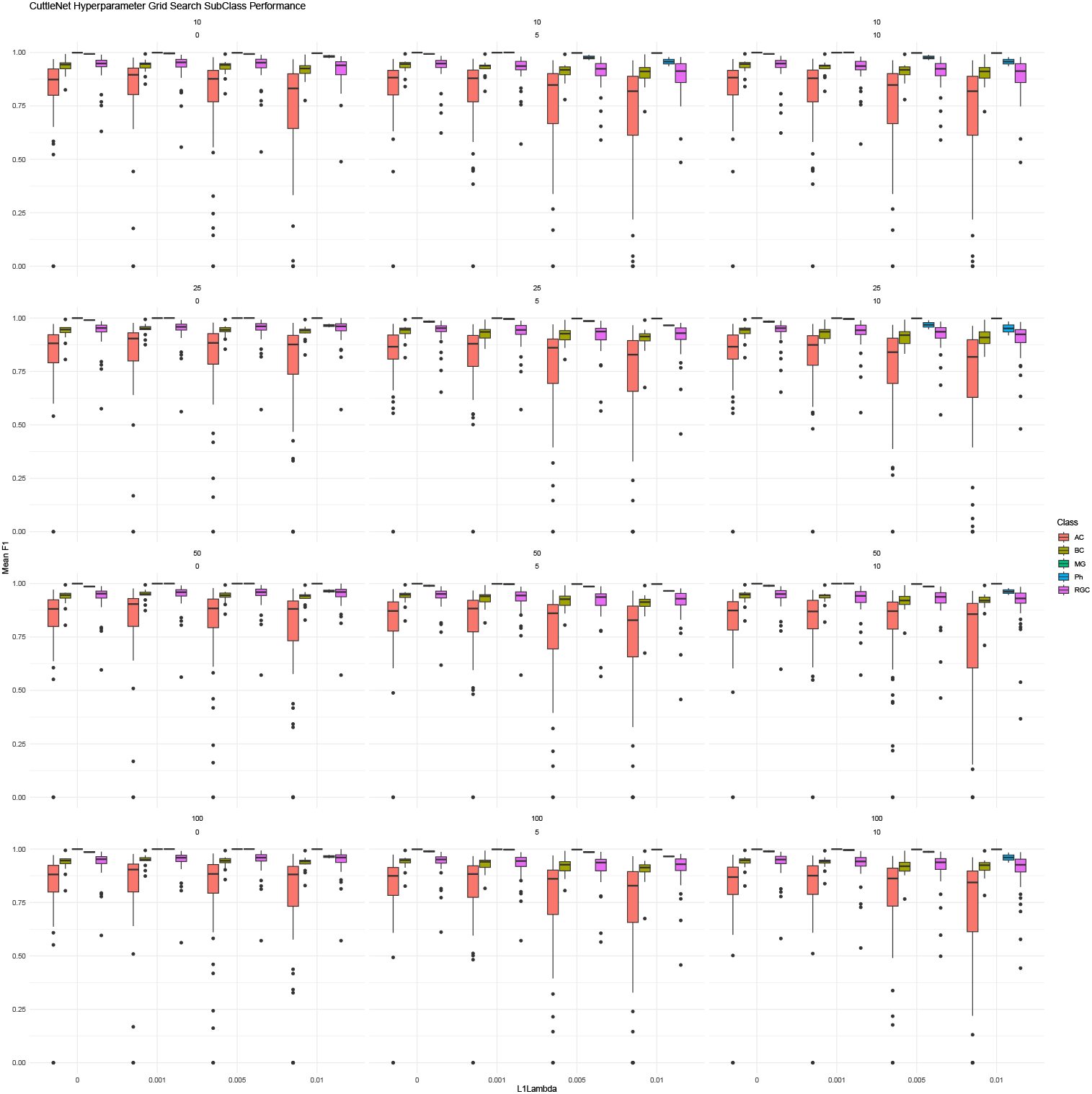
Grid search of CuttleNet with various hyperparameters that were explored. Panels are subset by Number of Epochs from top to bottom, Early Stopping parameter left to right. Colors show classes, box plots show F1 score of the subclasses, and X axis shows L1 Regularization parameter.

**Figure A.9:**
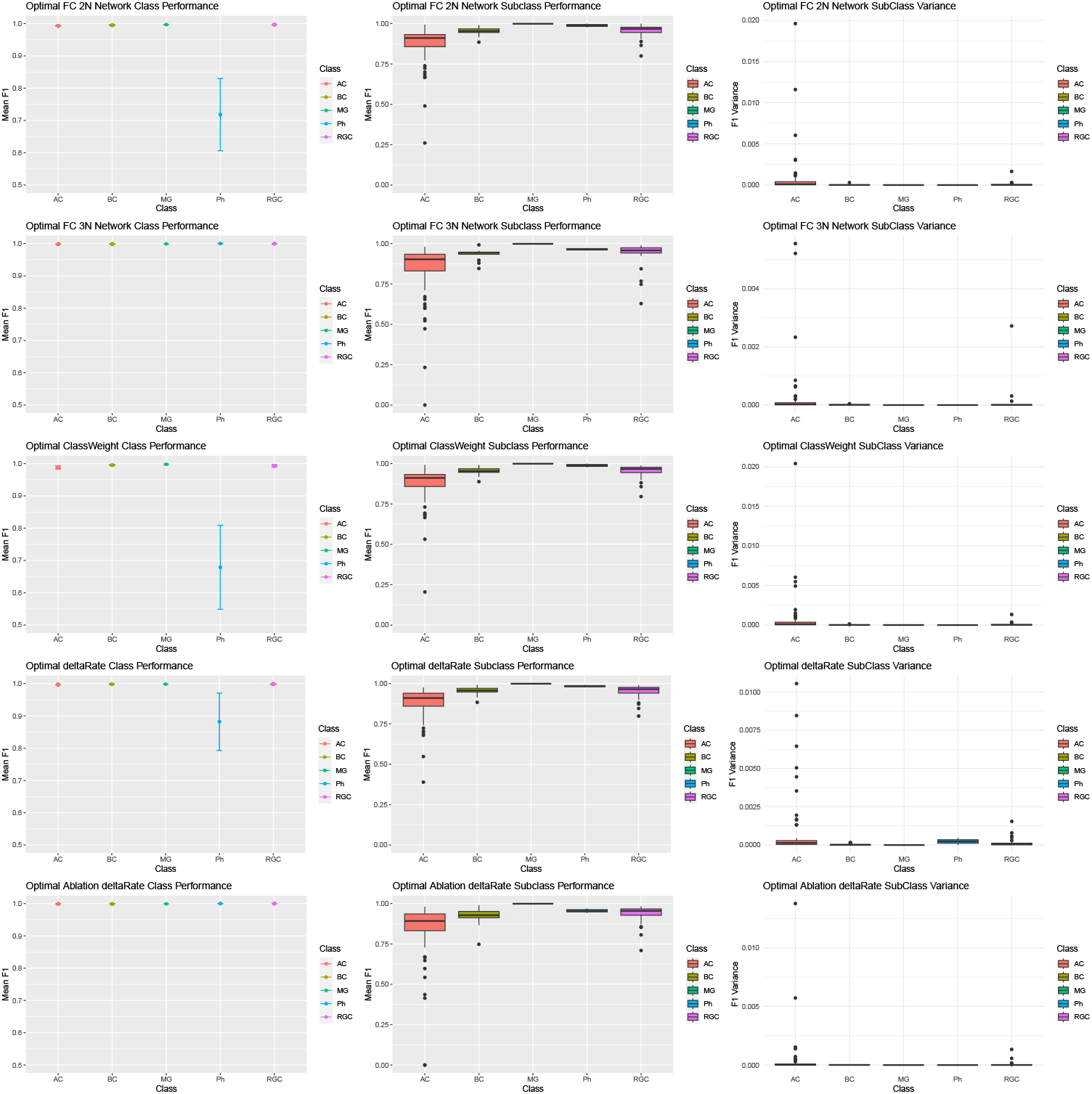
Optimal versions of simple NN vs more complicated variants considered but not used for GraSP. Here, each architecture evaluated after performing appropriate grid searches. Each experimental condition in the search included five replicates of training and evaluating the respective model; an 80/20 train/test split was done in all cases with the metrics evaluated here coming only from the testing dataset.

### A.5 Impact Statement

This paper presents an application which will advance the field spatial biology. Neither GraSP nor CuttleNet offer opportunities for abuse or negative societal consequence beyond historic ML methods.

#### NeurIPS Paper Checklist

##### 1. Claims

Question: Do the main claims made in the abstract and introduction accurately reflect the paper’s contributions and scope?

Answer: [Yes]

Justification: The abstract describes briefly the GraSP method and comparisons that were done to validate it, as well as the CuttleNet architecture and benchmarking it. These are the topics the manuscript covers in detail.

Guidelines:

- The answer NA means that the abstract and introduction do not include the claims made in the paper.
- The abstract and/or introduction should clearly state the claims made, including the contributions made in the paper and important assumptions and limitations. A No or NA answer to this question will not be perceived well by the reviewers.
- The claims made should match theoretical and experimental results, and reflect how much the results can be expected to generalize to other settings.
- It is fine to include aspirational goals as motivation as long as it is clear that these goals are not attained by the paper.

##### 2. Limitations

Question: Does the paper discuss the limitations of the work performed by the authors? Answer: [Yes]

Justification: We highlight that we have explored this ML method and architecture only in the limited application space of extracting gene predictors for spatial biology in the context of retinal data. We point out in the discussion that more work is needed to expand this work to see how generally applicable these ideas prove to be, and that despite our significant performance relative to benchmarks we have not demonstrated general superiority. Due to space constraints we did not have space available for a separate Limitations section.

Guidelines:

- The answer NA means that the paper has no limitation while the answer No means that the paper has limitations, but those are not discussed in the paper.
- The authors are encouraged to create a separate “Limitations” section in their paper.
- The paper should point out any strong assumptions and how robust the results are to violations of these assumptions (e.g., independence assumptions, noiseless settings, model well-specification, asymptotic approximations only holding locally). The authors should reflect on how these assumptions might be violated in practice and what the implications would be.
- The authors should reflect on the scope of the claims made, e.g., if the approach was only tested on a few datasets or with a few runs. In general, empirical results often depend on implicit assumptions, which should be articulated.
- The authors should reflect on the factors that influence the performance of the approach. For example, a facial recognition algorithm may perform poorly when image resolution is low or images are taken in low lighting. Or a speech-to-text system might not be used reliably to provide closed captions for online lectures because it fails to handle technical jargon.
- The authors should discuss the computational efficiency of the proposed algorithms and how they scale with dataset size.
- If applicable, the authors should discuss possible limitations of their approach to address problems of privacy and fairness.
- While the authors might fear that complete honesty about limitations might be used by reviewers as grounds for rejection, a worse outcome might be that reviewers discover limitations that aren’t acknowledged in the paper. The authors should use their best judgment and recognize that individual actions in favor of transparency play an important role in developing norms that preserve the integrity of the community. Reviewers will be specifically instructed to not penalize honesty concerning limitations.

##### 3. Theory Assumptions and Proofs

Question: For each theoretical result, does the paper provide the full set of assumptions and a complete (and correct) proof?

Answer: [NA]

Justification: We do not make theoretical claims, all of our claims are empirical. Guidelines:

- The answer NA means that the paper does not include theoretical results.
- All the theorems, formulas, and proofs in the paper should be numbered and cross-referenced.
- All assumptions should be clearly stated or referenced in the statement of any theorems.
- The proofs can either appear in the main paper or the supplemental material, but if they appear in the supplemental material, the authors are encouraged to provide a short proof sketch to provide intuition.
- Inversely, any informal proof provided in the core of the paper should be complemented by formal proofs provided in appendix or supplemental material.
- Theorems and Lemmas that the proof relies upon should be properly referenced.

##### 4. Experimental Result Reproducibility

Question: Does the paper fully disclose all the information needed to reproduce the main experimental results of the paper to the extent that it affects the main claims and/or conclusions of the paper (regardless of whether the code and data are provided or not)?

Answer: [Yes]

Justification: Yes all code is provided as are the relevant open access datasets. Moreover, within the space constraints we fully disclosed how and what we did for each experiment and analysis.

Guidelines:

- The answer NA means that the paper does not include experiments.
- If the paper includes experiments, a No answer to this question will not be perceived well by the reviewers: Making the paper reproducible is important, regardless of whether the code and data are provided or not.
- If the contribution is a dataset and/or model, the authors should describe the steps taken to make their results reproducible or verifiable.
- Depending on the contribution, reprehensibility can be accomplished in various ways. For example, if the contribution is a novel architecture, describing the architecture fully might suffice, or if the contribution is a specific model and empirical evaluation, it may be necessary to either make it possible for others to replicate the model with the same dataset, or provide access to the model. In general. releasing code and data is often one good way to accomplish this, but reproducibility can also be provided via detailed instructions for how to replicate the results, access to a hosted model (e.g., in the case of a large language model), releasing of a model checkpoint, or other means that are appropriate to the research performed.
- While NeurIPS does not require releasing code, the conference does require all submissions to provide some reasonable avenue for reproducibility, which may depend on the nature of the contribution. For example
  a. If the contribution is primarily a new algorithm, the paper should make it clear how to reproduce that algorithm.
  b. If the contribution is primarily a new model architecture, the paper should describe the architecture clearly and fully.
  c. If the contribution is a new model (e.g., a large language model), then there should either be a way to access this model for reproducing the results or a way to reproduce the model (e.g., with an open-source dataset or instructions for how to construct the dataset).
  d. We recognize that reproducibility may be tricky in some cases, in which case authors are welcome to describe the particular way they provide for reproducibility.

In the case of closed-source models, it may be that access to the model is limited in some way (e.g., to registered users), but it should be possible for other researchers to have some path to reproducing or verifying the results.

##### 5. Open access to data and code

Question: Does the paper provide open access to the data and code, with sufficient instructions to faithfully reproduce the main experimental results, as described in supplemental material?

Answer: [Yes]

Justification: All code is provided in an anonymyzed github repository, and all datasets are avialble on the internet and clearly listed.

Guidelines:

- The answer NA means that paper does not include experiments requiring code.
- Please see the NeurIPS code and data submission guidelines (https://nips.cc/public/guides/CodeSubmissionPolicy) for more details.
- While we encourage the release of code and data, we understand that this might not be possible, so “No” is an acceptable answer. Papers cannot be rejected simply for not including code, unless this is central to the contribution (e.g., for a new open-source benchmark).
- The instructions should contain the exact command and environment needed to run to reproduce the results. See the NeurIPS code and data submission guidelines (https://nips.cc/public/guides/CodeSubmissionPolicy) for more details.
- The authors should provide instructions on data access and preparation, including how to access the raw data, preprocessed data, intermediate data, and generated data, etc.
- The authors should provide scripts to reproduce all experimental results for the new proposed method and baselines. If only a subset of experiments are reproducible, they should state which ones are omitted from the script and why.
- At submission time, to preserve anonymity, the authors should release anonymized versions (if applicable).
- Providing as much information as possible in supplemental material (appended to the paper) is recommended, but including URLs to data and code is permitted.

##### 6. Experimental Setting/Details

Question: Does the paper specify all the training and test details (e.g., data splits, hyperparameters, how they were chosen, type of optimizer, etc.) necessary to understand the results?

Answer: [Yes]

Justification: We listed in as much detail narratively as we could all hyper-parameter choices. Moreover, the code is provided should any clarification be necessary.

Guidelines:

- The answer NA means that the paper does not include experiments.
- The experimental setting should be presented in the core of the paper to a level of detail that is necessary to appreciate the results and make sense of them.
- The full details can be provided either with the code, in appendix, or as supplemental material.

##### 7. Experiment Statistical Significance

Question: Does the paper report error bars suitably and correctly defined or other appropriate information about the statistical significance of the experiments?

Answer: [Yes]

Justification: Standard Boxplots are used for all plots so full distributions are visible.All statistics are listed and tests explained.

Guidelines:

- The answer NA means that the paper does not include experiments.
- The authors should answer “Yes” if the results are accompanied by error bars, confidence intervals, or statistical significance tests, at least for the experiments that support the main claims of the paper.
- The factors of variability that the error bars are capturing should be clearly stated (for example, train/test split, initialization, random drawing of some parameter, or overall run with given experimental conditions).
- The method for calculating the error bars should be explained (closed form formula, call to a library function, bootstrap, etc.)
- The assumptions made should be given (e.g., Normally distributed errors).
- It should be clear whether the error bar is the standard deviation or the standard error of the mean.
- It is OK to report 1-sigma error bars, but one should state it. The authors should preferably report a 2-sigma error bar than state that they have a 96% CI, if the hypothesis of Normality of errors is not verified.
- For asymmetric distributions, the authors should be careful not to show in tables or figures symmetric error bars that would yield results that are out of range (e.g. negative error rates).
- If error bars are reported in tables or plots, The authors should explain in the text how they were calculated and reference the corresponding figures or tables in the text.

##### 8. Experiments Compute Resources

Question: For each experiment, does the paper provide sufficient information on the computer resources (type of compute workers, memory, time of execution) needed to reproduce the experiments?

Answer: [Yes]

Justification: Hardware is listed, execution time is listed for the traditional ML methods and GraSP briefly. We briefly reference some of the many other exploratory experiments we performed which were computationally intensive but did not converge (genetic evolutionary algorithms, GBM), these experiments occurred over the course of several years and were not repeated with the present more focused evaluation of our final algorithm. Finally, the methods we are presenting are simply not computationally intensive as we are mostly training neural networks with fewer than 5 layers as reviewers in this audience should appreciate.

Guidelines:

- The answer NA means that the paper does not include experiments.
- The paper should indicate the type of compute workers CPU or GPU, internal cluster, or cloud provider, including relevant memory and storage.
- The paper should provide the amount of compute required for each of the individual experimental runs as well as estimate the total compute.
- The paper should disclose whether the full research project required more compute than the experiments reported in the paper (e.g., preliminary or failed experiments that didn’t make it into the paper).

##### 9. Code Of Ethics

Question: Does the research conducted in the paper conform, in every respect, with the NeurIPS Code of Ethics https://neurips.cc/public/EthicsGuidelines?

Answer: [Yes]

Justification: We do not violate these code of ethics.

Guidelines:

- The answer NA means that the authors have not reviewed the NeurIPS Code of Ethics.
- If the authors answer No, they should explain the special circumstances that require a deviation from the Code of Ethics.
- The authors should make sure to preserve anonymity (e.g., if there is a special consideration due to laws or regulations in their jurisdiction).

##### 10. Broader Impacts

Question: Does the paper discuss both potential positive societal impacts and negative societal impacts of the work performed?

Answer: [NA]

Justification: We have an impacts statement in the appendix, but ultimately we do not anticipate any material impacts of our work.

Guidelines:

- The answer NA means that there is no societal impact of the work performed.
- If the authors answer NA or No, they should explain why their work has no societal impact or why the paper does not address societal impact.
- Examples of negative societal impacts include potential malicious or unintended uses (e.g., disinformation, generating fake profiles, surveillance), fairness considerations (e.g., deployment of technologies that could make decisions that unfairly impact specific groups), privacy considerations, and security considerations.
- The conference expects that many papers will be foundational research and not tied to particular applications, let alone deployments. However, if there is a direct path to any negative applications, the authors should point it out. For example, it is legitimate to point out that an improvement in the quality of generative models could be used to generate deepfakes for disinformation. On the other hand, it is not needed to point out that a generic algorithm for optimizing neural networks could enable people to train models that generate Deepfakes faster.
- The authors should consider possible harms that could arise when the technology is being used as intended and functioning correctly, harms that could arise when the technology is being used as intended but gives incorrect results, and harms following from (intentional or unintentional) misuse of the technology.
- If there are negative societal impacts, the authors could also discuss possible mitigation strategies (e.g., gated release of models, providing defenses in addition to attacks, mechanisms for monitoring misuse, mechanisms to monitor how a system learns from feedback over time, improving the efficiency and accessibility of ML).

##### 11. Safeguards

Question: Does the paper describe safeguards that have been put in place for responsible release of data or models that have a high risk for misuse (e.g., pretrained language models, image generators, or scraped datasets)?

Answer: [NA]

Justification: We do not foresee misuse of our methods.

Guidelines:

- The answer NA means that the paper poses no such risks.
- Released models that have a high risk for misuse or dual-use should be released with necessary safeguards to allow for controlled use of the model, for example by requiring that users adhere to usage guidelines or restrictions to access the model or implementing safety filters.
- Datasets that have been scraped from the Internet could pose safety risks. The authors should describe how they avoided releasing unsafe images.
- We recognize that providing effective safeguards is challenging, and many papers do not require this, but we encourage authors to take this into account and make a best faith effort.

##### 12. Licenses for existing assets

Question: Are the creators or original owners of assets (e.g., code, data, models), used in the paper, properly credited and are the license and terms of use explicitly mentioned and properly respected?

Answer: [Yes]

Justification: The R and python libraries we use are clearly listed, as are the datasets. All other assets belong to the authors.

Guidelines:

- The answer NA means that the paper does not use existing assets.
- The authors should cite the original paper that produced the code package or dataset.
- The authors should state which version of the asset is used and, if possible, include a URL.
- The name of the license (e.g., CC-BY 4.0) should be included for each asset.
- For scraped data from a particular source (e.g., website), the copyright and terms of service of that source should be provided.
- If assets are released, the license, copyright information, and terms of use in the package should be provided. For popular datasets, paperswithcode.com/datasets has curated licenses for some datasets. Their licensing guide can help determine the license of a dataset.
- For existing datasets that are re-packaged, both the original license and the license of the derived asset (if it has changed) should be provided.
- If this information is not available online, the authors are encouraged to reach out to the asset’s creators.

##### 13. New Assets

Question: Are new assets introduced in the paper well documented and is the documentation provided alongside the assets?

Answer: [No]

Justification: We provided details and code for others to reproduce our gene target list, however, we are actively pursuing this biological research and plan to release the target list with that publication in a different venue focused on the biologic results. All models can be regenerated from the provided code.

Guidelines:

- The answer NA means that the paper does not release new assets.
- Researchers should communicate the details of the dataset/code/model as part of their submissions via structured templates. This includes details about training, license, limitations, etc.
- The paper should discuss whether and how consent was obtained from people whose asset is used.
- At submission time, remember to anonymize your assets (if applicable). You can either create an anonymized URL or include an anonymized zip file.

##### 14. Crowdsourcing and Research with Human Subjects

Question: For crowdsourcing experiments and research with human subjects, does the paper include the full text of instructions given to participants and screenshots, if applicable, as well as details about compensation (if any)?

Answer: [NA]

Justification: The only people involved were the authors.

Guidelines:

- The answer NA means that the paper does not involve crowdsourcing nor research with human subjects.
- Including this information in the supplemental material is fine, but if the main contribution of the paper involves human subjects, then as much detail as possible should be included in the main paper.
- According to the NeurIPS Code of Ethics, workers involved in data collection, curation, or other labor should be paid at least the minimum wage in the country of the data collector.

##### 15. Institutional Review Board (IRB) Approvals or Equivalent for Research with Human Subjects

Question: Does the paper describe potential risks incurred by study participants, whether such risks were disclosed to the subjects, and whether Institutional Review Board (IRB) approvals (or an equivalent approval/review based on the requirements of your country or institution) were obtained?

Answer: [NA]

Justification: No human research was conducted.

Guidelines:

- The answer NA means that the paper does not involve crowdsourcing nor research with human subjects.
- Depending on the country in which research is conducted, IRB approval (or equivalent) may be required for any human subjects research. If you obtained IRB approval, you should clearly state this in the paper.
- We recognize that the procedures for this may vary significantly between institutions and locations, and we expect authors to adhere to the NeurIPS Code of Ethics and the guidelines for their institution.
- For initial submissions, do not include any information that would break anonymity (if applicable), such as the institution conducting the review.

## Footnotes

2 For example, the alphaRGCs mediate high acuity vision and can be easily distinguished from other subclasses based on a small set of genes. Due to the relative simplicity with which they could be classified, spatial maps of these cells were produced [5] which directly explained hunting behaviors observed in mice [6].

3 The post-hoc testing discussed below revealed no significant differences between the specific thresholds.

4 See appendix for more specific observations from this grid search.

